# Chromosome polymorphisms track trans-Atlantic divergence, admixture and adaptive evolution in salmon

**DOI:** 10.1101/351338

**Authors:** Sarah J. Lehnert, Paul Bentzen, Tony Kess, Sigbjørn Lien, John B. Horne, Marie Clément, Ian R. Bradbury

## Abstract

Pleistocene glaciations drove repeated range contractions and expansions shaping contemporary intraspecific diversity. Atlantic salmon (*Salmo salar*) from the western and eastern Atlantic range diverged >600K YBP, with each clade isolated in independent southern refugia during glacial maxima, driving trans-Atlantic genomic and karyotypic differences. Here, we investigate genomic consequences of glacial isolation and transAtlantic secondary contact using a 220K single nucleotide polymorphism (SNP) array genotyped in 80 North American and European populations. Throughout North America, we identified large inter-individual variation and discrete linkage blocks within and between chromosomes with known rearrangements: Ssa01/Ssa23 translocation and Ssa08/Ssa29 fusion. Spatial genetic analyses suggest independence of rearrangements, with Ssa01/Ssa23 showing high European introgression (>50%) in northern populations indicative of post-glacial trans-Atlantic secondary contact, contrasting low European ancestry genome-wide (3%). Ssa08/Ssa29 showed greater intra-population diversity suggesting a derived chromosome fusion polymorphism within North America. Evidence of selection on both regions suggests adaptive variation associated with karyotypes. Our study highlights how Pleistocene glaciations can drive large-scale intraspecific variation in genomic architecture of northern species.

## Introduction

The Pleistocene glaciations resulted in global climatic shifts driving repeated periods of isolation and range expansions that have shaped contemporary biodiversity in North America and Europe (1, 2). Glaciers covered much of northern regions, limiting movement and connectivity within species and forcing populations into glacial refugia for prolonged periods of isolation (~100,000 years) (1–3). When ice sheets retreated following glacial maxima, colonization of newly available habitats enabled secondary contact between allopatric lineages from different refugia (1, 3, 4). Secondary contact between divergent lineages can have significant evolutionary consequences (5–10), and the genetic legacy of these post-glacial admixture and range expansion events are present in modern genomes of many northern species (1, 3, 11–13).

Atlantic salmon (*Salmo salar*) is an anadromous species of significant ecological, social, and economic value across its range (14). Populations of Atlantic salmon span both sides of the North Atlantic Ocean where populations are highly structured across multiple spatial scales with the deepest split occurring between continents (15, 16). Salmon in the western and eastern Atlantic are thought to have diverged >600K YBP (16–18). These clades occupied different refugia during glacial maxima, accruing many genetic differences including differences in chromosome number and structure (19) resulting in genetic incompatibilities (20). Nonetheless, despite lengthy periods of isolation, genetic data provide evidence of trans-Atlantic secondary contact in Europe and North America near the end of last glacial maximum (~10K YBP) when glaciers retreated and salmon recolonized much of their contemporary range (16, 18, 21–23). The full extent of trans-Atlantic secondary contact across much of the range remains largely unknown; however, within North America, ‘European’ mitochondrial haplotypes have been documented in Newfoundland, Quebec, and Labrador (16, 17, 24) and genome-wide evidence suggests that signatures of secondary contact are not limited to the mitochondrial genome (18, 21).

The genomic consequences of these secondary contact events can depend on genomic architecture where differences in chromosomal structure (*i.e.*, inversions and/or translocations) can act as post-zygotic barriers to gene flow (25) leading to heterogeneous genomic introgression (26, 27). However, admixture can also introduce new chromosomal variants already shaped by selection (6, 25), such as inversions in the Apple maggot (*Rhagoletis pomonella*) (9) or Robertsonian rearrangements in the house mouse (*Mus musculus domesticus*) (28, 29), that can facilitate adaptive divergence, range expansions, and ultimately speciation (8, 30, 31). Limited karyotype data suggest that in North American Atlantic salmon, structural rearrangements have reduced the number of chromosome pairs from 29 to 27, including two chromosome fusions (Ssa08/Ssa29 and Ssa26/Ssa28) and one translocation with a fission (Ssa01p/Ssa23 and Ssa01q), although heterogeneity in these rearrangements has been reported (19). We hypothesize that genomic regions associated with rearrangements may carry evidence of trans-Atlantic secondary contact events as these regions should be characterized by reduced recombination in hetero-karyotype individuals allowing signatures of contact to persist over time.

To explore how the Pleistocene glaciation has shaped the modern genome of Atlantic salmon, we use high-density genome-wide single nucleotide polymorphisms (SNPs) to identify large chromosomal regions reflecting signals of historical transAtlantic secondary contact in North American Atlantic salmon. Our study highlights how periods of glacial isolation, secondary contact, and genomic architecture contribute to heterogeneous genomic introgression that drives intraspecific diversity.

## Results

### Detection of individual-based differences in genomic architecture

Atlantic salmon (n = 1459) from 80 populations spanning Norway and Canada (Fig. 1AB; Table S1) were genotyped using a 220,000 target, bi-allelic SNP Affymetrix Axiom array developed for Atlantic salmon. To explore individual-based differences in genomic architecture across multiple spatial scales, we used the R package *pcadapt* (32) which detects candidate genomic regions of differentiation using principal component (PC) analysis with large datasets that can contain admixed individuals and hierarchical population structure. Within North America, genetic variation separated populations by geographic regions along the first two PC axes (Fig. 1C), while an increasing number of PC axes tested (K = 25 to 50; see Supplement Fig. S1, S2) identified large genomic regions (>5 Mbp) associated with individual variation on four chromosomes (based on the European genome) with known structural rearrangements in Atlantic salmon (19) (Fig. 1D; Table S2). We consider these four genomic regions (Ssa01, Ssa08, Ssa23, and Ssa29) to be “outlier blocks” as each contained at least 200 SNPs that were statistical outliers (adjusted *p*-value [*q*-value] < 0.05) (Table S2).

**Fig. 1.**
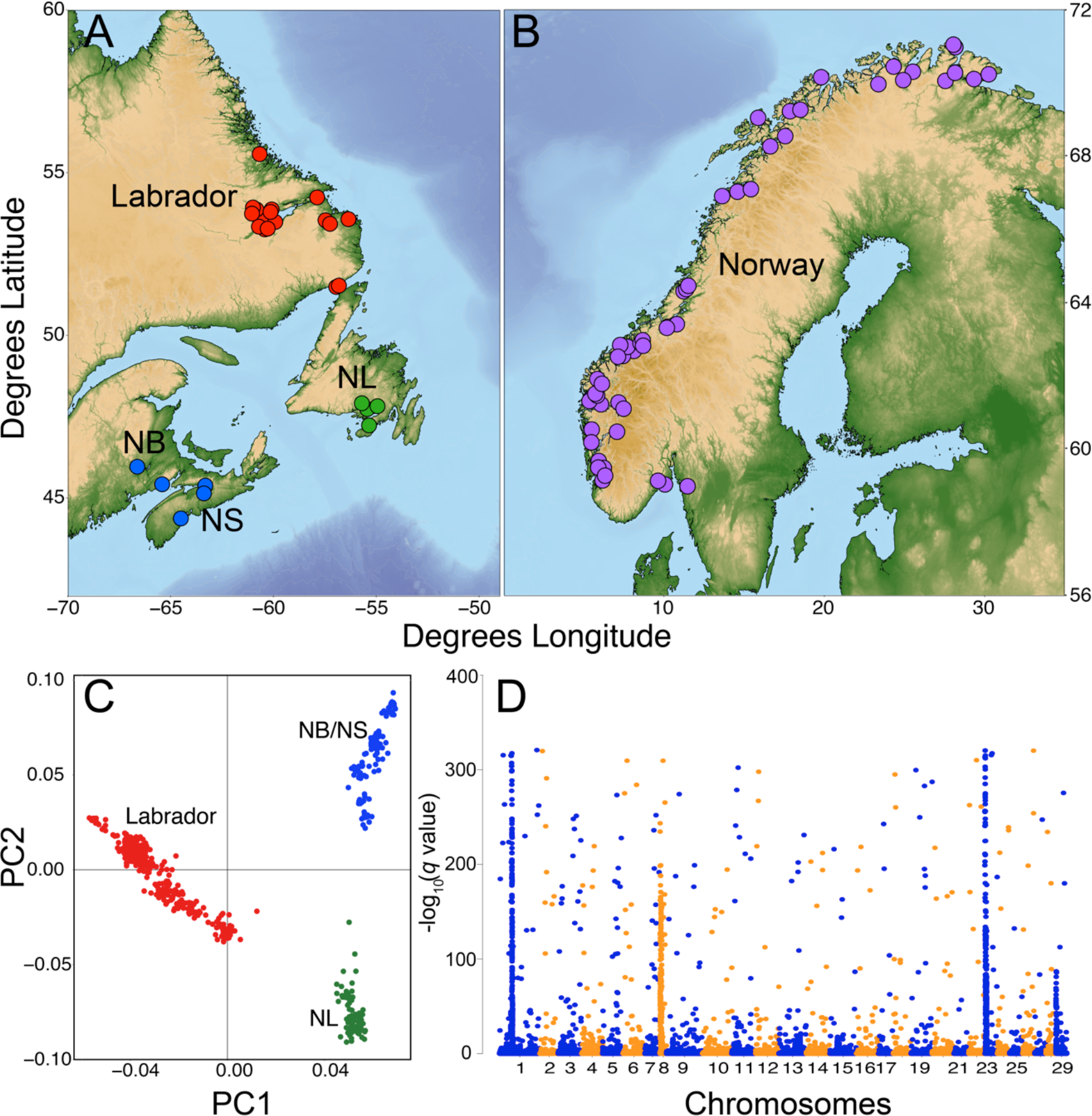
Sampling sites of Atlantic salmon (*S. salar*) across the North Atlantic Ocean and the detection of individual-based differences in genomic architecture within North America. Maps of sampling sites in **(A)** Canada and **(B)** Norway with sites coloured by geographic region. **(C)** Regional level population structure across Canadian populations based on the first two principal components (PC) axes from *pcadapt* (32) using 104,119 single nucleotide polymorphisms (SNPs). **(D)** Manhattan plot of adjusted *p*-values (*q*-values) showing genomic regions of significant inter-individual variation based on the retention of 40 PC axes in *pcadapt*.

### Linkage disequilibrium of outlier blocks

A heatmap of linkage disequilibrium (LD) between all outlier SNPs (*q*-value < 0.05) from Ssa01, Ssa08, Ssa23, and Ssa29 revealed regions of high LD consistent with known trans-Atlantic rearrangements, including high LD within and between chromosomes with a translocation (Ssa01 and Ssa23) and chromosomes with a fusion (Ssa08 and Ssa29) in North America (Fig. 2A). These high LD regions were only present in Canada and not in Norway (Fig. 2A). Using all SNPs (MAF > 0.05) from these pairs of chromosomes for Canadian populations, we identified a region of high LD between Ssa01 and Ssa23 SNPs (Fig. 2B) and another between Ssa08 and Ssa29 SNPs (Fig. 2C). These regions were further examined within each chromosome where high LD regions (measured as mean pairwise LD) generally corresponded to outlier blocks identified in *pcadapt* (Fig. 3 and Table S2) and high LD was not present in Norway. Given high LD between blocks from different chromosomes, outlier SNPs were combined and analyzed together (Ssa01/Ssa23 and Ssa08/Ssa29) for remaining analyses.

**Fig. 2.**
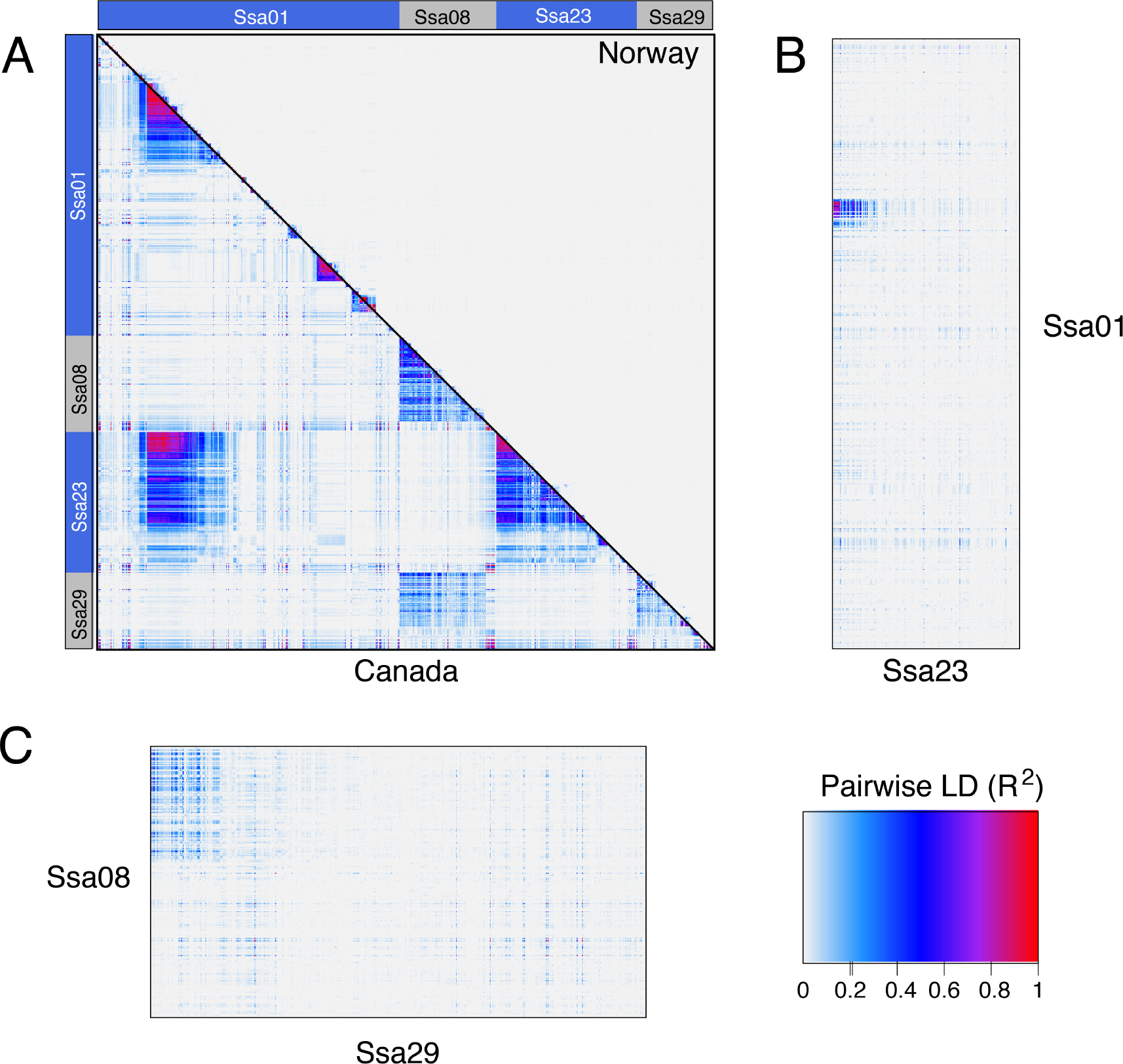
Linkage disequilibrium (LD) between outliers from chromosomes with known rearrangements in Atlantic salmon (*S. salar*). **(A)** Heatmap of pairwise LD calculated as R^2^ for outlier loci associated with individual-based variation on four chromosomes (Ssa01, 08, 23, and 29) across Canadian populations using *pcadapt* (32). Chromosomes are indicated by gray and blue bars along the plot edges. The lower matrix represents pairwise LD for all Canadian samples and the upper matrix represents pairwise LD for all Norwegian samples. **(B, C)** Heatmaps of pairwise LD between chromosomes across Canadian samples, where **(B)** represents all loci (filtered for minor allele frequency >0.05) on Ssa01 and Ssa23 and **(C)** on Ssa08 and Ssa29. Note that heatmaps in **(B)** and **(C)** are not scaled relative to each other.

**Fig. 3.**
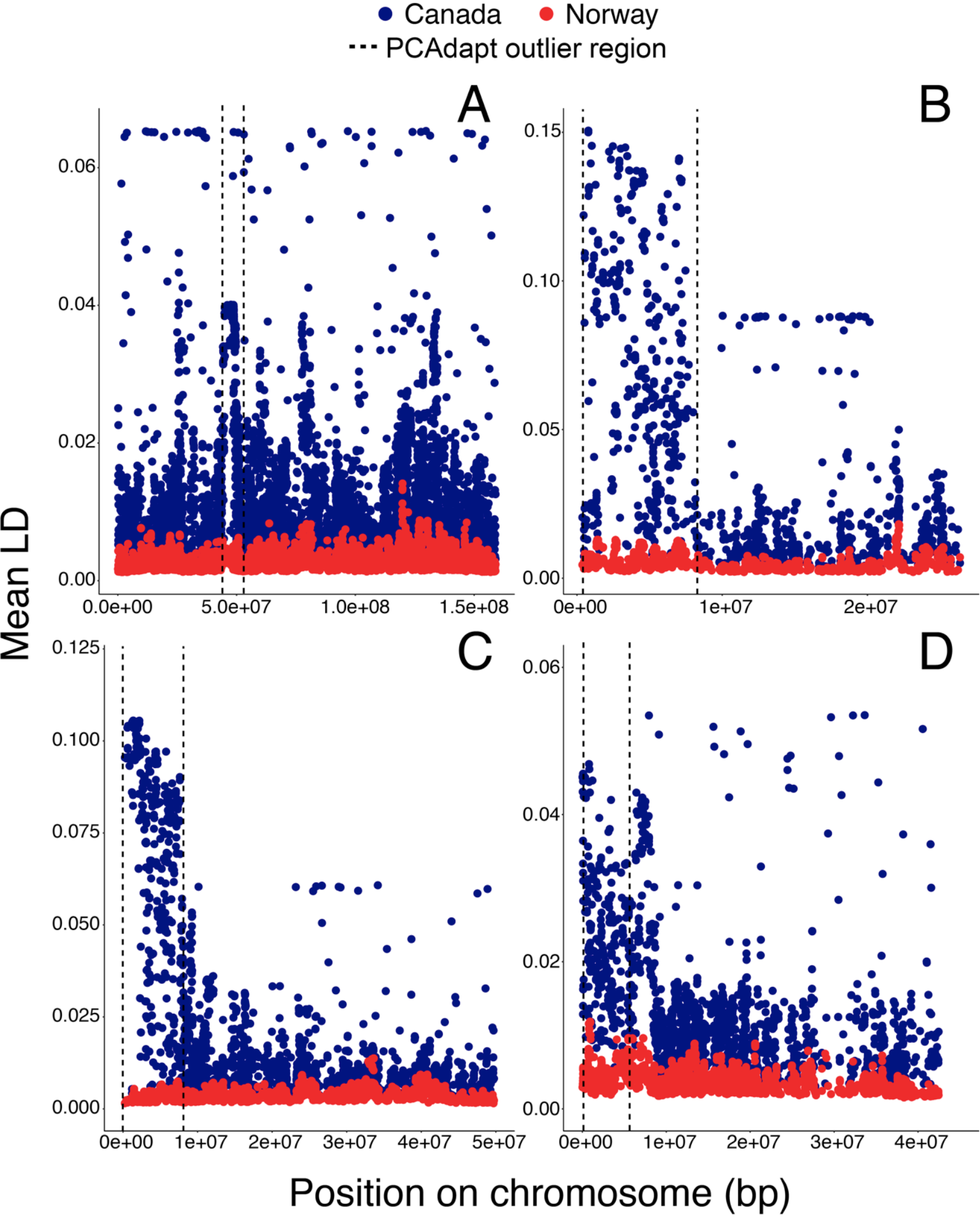
Mean pairwise linkage disequilibrium (LD) across known rearranged chromosomes within Norwegian (red) and Canadian (blue) Atlantic salmon (*S. salar*). Mean LD was calculated for each locus as the mean pairwise R^2^ across all loci (with minor allele frequency >0.05 in Canada) on the chromosome. Panels represent **(A)** Ssa01, **(B)** Ssa08, **(C)** Ssa23, and **(D)** Ssa29. Positions are based on the European Atlantic salmon genome and dashed lines represent outlier block regions detected in *pcadapt*.

### Population and individual level genetic structure of outlier blocks

Bayesian clustering for both outlier datasets were performed in STRUCTURE (33). Using the ΔK statistic (34), the optimal number of genetic clusters was identified as K = 2 for both datasets (see Fig. S3; note some support for K = 3 for the Ssa08/Ssa29 dataset, see below) and Norwegian sites were almost entirely comprised of a single cluster (European cluster) but Canadian sites varied in their proportion of membership to the two (European and North American) clusters (Fig. 4). For Ssa01/Ssa23, the proportion of membership to the European cluster increased with latitude in Canada while many southern sites were fixed for the North American genetic cluster (Fig. 4AB). Populations in Labrador, primarily within the Lake Melville system (a 3,069 km^2^ marine embayment), showed a greater proportion of membership to the European cluster than other sites (Fig. 4AB). Clustering patterns for Ssa08/Ssa29 and Ssa01/Ssa23 were not consistent across Canadian populations (Fig. 4CD). For Ssa08/Ssa29, all populations in Canada were assigned membership to both the European and North American clusters, with the North American genetic cluster not reaching fixation in any population (Fig. 4C). However, for Ssa08/Ssa29 there was also some support for K = 3 which separated Europe from North America, suggesting two genetic groups in North America and a distinct cluster in Europe (see Fig. S3 and S4). Using generalized linear models, population allele frequencies across latitude were examined within outlier blocks and more SNPs from the Ssa01/Ssa23 blocks demonstrated latitudinal clines (Fig. S5A) whereas SNPs from Ssa08/Ssa29 (Fig. S5B) were more variable across latitude.

**Fig. 4.**
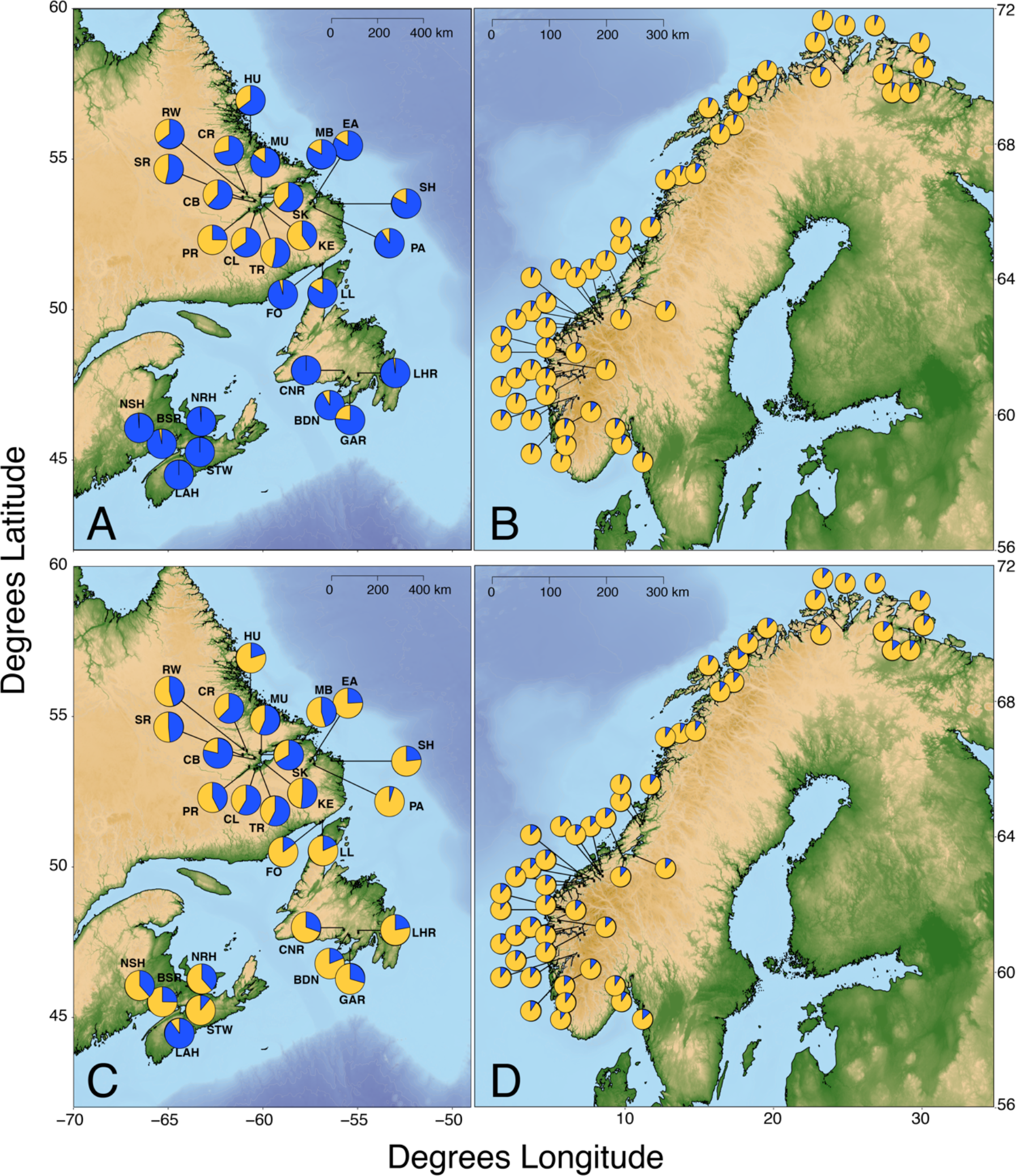
Bayesian clustering of genomic outlier blocks across populations of Atlantic salmon (*S. salar*). Maps of population structure for **(A-B)** outlier single nucleotide polymorphisms (SNPs) on outlier blocks of Ssa01 and Ssa23 and **(C-D)** Ssa08 and Ssa29 for Atlantic salmon across 80 sampling locations in **(A, C)** Canada and **(B, D)** Norway. Panels show the results of Bayesian clustering analysis with the proportion membership assigned to two genetic clusters for each population.

Results of Bayesian clustering suggest different evolutionary histories for these two rearrangements, thus to further investigate these potential differences, genetic relationships in outlier blocks were examined among individuals using principal component analysis (PCA) and individual-based neighbour-joining (NJ) trees. First, a panel of genome-wide non-outlier loci (1500 randomly selected SNPs with *pcadapt q-* value > 0.05 and excluding rearranged chromosomes) showed a clear trans-Atlantic split for both the PCA and NJ tree (Fig. 5AB). Conversely, when examining SNPs from outlier blocks on Ssa01/Ssa23, reduced trans-Atlantic discontinuity was visible, with individual branches generally corresponding to variation in Bayesian cluster membership (Fig. 5CD). Some individuals from Labrador (primarily within the Lake Melville system) grouped with Norwegian individuals in the NJ tree (Fig. 5D) and another intermediate cluster of individuals was present between European and the “pure North American” (based on STRUCTURE) individuals (Fig. 5D) suggesting evidence of trans-Atlantic secondary contact. Clustering for Ssa08/29 showed different patterns with many individuals from all regions forming a group close but separate from Norwegian individuals (Fig. 5EF) suggesting an ancestral North American karyotype which is European-like (no Ssas08/Ssa29 fusion). Two additional clusters on the NJ tree and PCA may reflect individuals heterozygous or homozygous for derived North American variation associated with the chromosomal fusion (see Fig. 5EF).

**Fig. 5.**
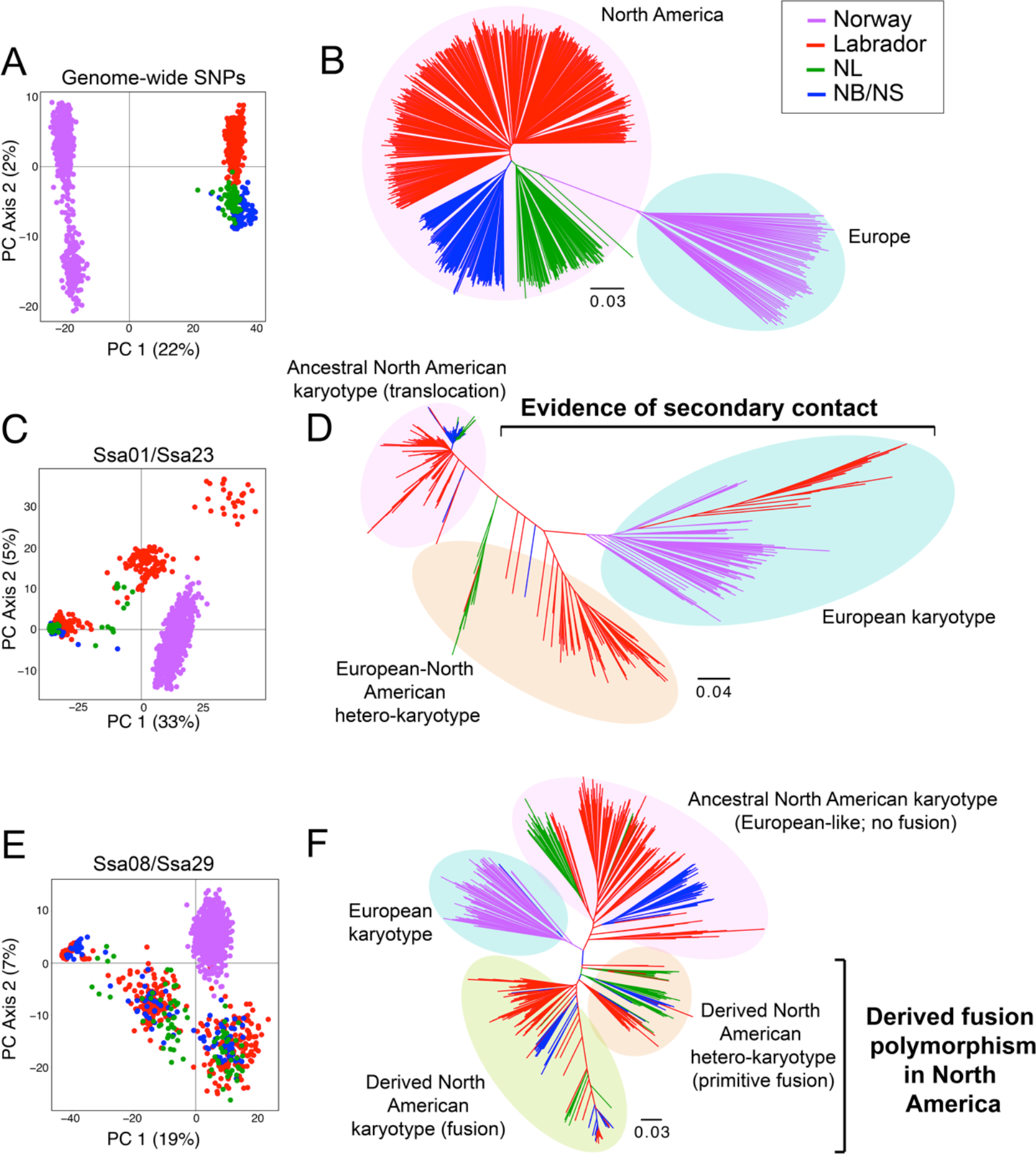
Genetic relationships for genome-wide non-outlier SNPs and genomic outlier blocks across individual Atlantic salmon (*S. salar*). Principal component analyses (PCA) and neighbour-joining (NJ) trees based on Cavalli-Sforza and Edwards (71) chord distances for individual salmon from Norway and Canada. **(A)** PC axis 1 and 2 and **(B)** NJ tree for a random set of 1500 single nucleotide polymorphisms (SNPs) that were nonsignificant in our *pcadapt* analysis and not located on the chromosomes with rearrangements. PCA and NJ tree results for significant outlier loci from **(C, D)** outlier blocks on Ssa01/Ssa23 and **(E, F)** outlier blocks on Ssa08/Ssa29. Branches and points are coloured by geographic region as in Fig. 1. All individuals were used for the PCA, whereas for the NJ tree, all North American individuals were include but only a subset of representative Norwegian individuals were chosen from five arbitrarily selected populations across different geographic regions (Enningdalselva, Nausta, Argardsvassdraget, Elvegardselva, and Risfjordvassdraget). The inferred karyotypes and histories of rearrangements are indicated with circles and text on the NJ trees based on the results of Bayesian clustering, NJ tree, and PCA.

### Ancestry and selection along chromosomes

Ancestry was assigned along each chromosome (with genomic positions aligned to the European genome) using PCAdmix (35) for North American individuals based on Bayesian clustering results of outlier regions. Across chromosomes Ssa01 and Ssa23, individuals with evidence of potential European admixed (STRUCTURE Q-values < 0.9; n = 191) were assigned ancestry to either ‘pure’ Europe (Q <0.1) or ‘pure’ North America (Q > 0.9), and we found clear signals of European ancestry within both outlier regions in the majority of admixed individuals (Fig. 6AB). High levels of European ancestry (mean European ancestry was >50% in some SNP windows across samples) in these outlier regions contrasted the low genome-wide European ancestry calculated across all chromosomes (mean ± standard error: 3.30 ± 0.07%) (Fig. S6). Notably, one individual showed evidence of recent European ancestry (potential second generation backcross) and may reflect introgression from an aquaculture escapee or trans-Atlantic straying event (Fig. 6AB). Next, Tajima’s D values across Ssa01 and Ssa23 chromosomes were consistent with positive selection (Mann-Whitney U test) on the ‘pure’ North American type within the outlier region of Ssa01 (W = 550, p = 8.39e-09; Fig. 6E) and Ssa23 (W = 357, p = 0.0004; Fig. 6F). Evidence of positive selection in the Ssa01 region was also found when pure North American and individuals with strong evidence of secondary contact (Q < 0.1) were combined (W = 1511, p = 0.0008; Fig. 6E), potentially driven by the larger sample size of the pure North American group (n = 355) relative to the admixed (n = 30). Consistent with these results, outlier regions in pure North American individuals showed significant evidence of a positive selective sweep for Ssa01 (Mann-Whitney U test, W = 4796, p = 1.01e-07) and Ssa23 (W = 1106, p = 0.011), but not in admixed individuals (Ssa01: W = 3317.5, p = 0.2457; Ssa23: W = 902.5, p = 0.374) (see Supplement Fig. S7).

**Fig. 6.**
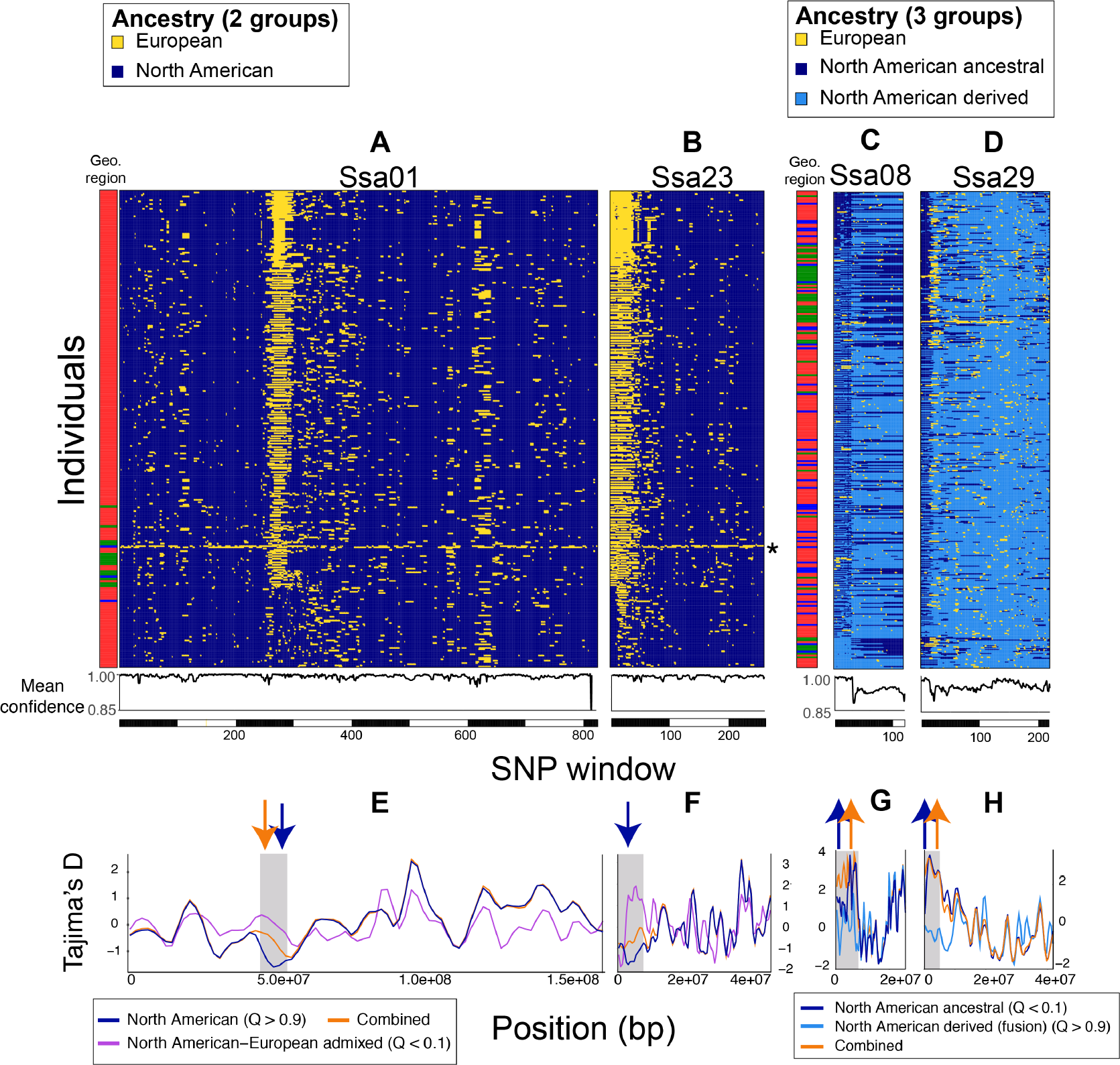
Ancestry and selection along rearranged chromosomes in North American Atlantic salmon (*S. salar*). Ancestry assigned for windows of 20 SNP along chromosomes **(A)** Ssa01, **(B)** Ssa23, **(C)** Ssa08, and **(D)** Ssa29 for North American salmon determined by PCAdmix (35) with pure and admixed groups separated based on STRUCTURE Q-values. For **(A,B)** Ssa01/Ssa23 ancestry was assigned to two groups including European (yellow) and North American (dark blue) for all individuals with evidence of European ancestry (Q-value < 0.9; n = 191). For **(C,D)** Ssa08/Ssa29 ancestry was assigned to three groups including European (yellow), North American ancestral (dark blue), and North American derived (light blue) for individuals with intermediate Q-values (0.1 < Q < 0.9; n = 250). Homologous chromosomes are shown together for each individual with individuals organized vertically by increasing Q-value (Ssa01/Ssa23: more to less European; Ssa08/Ssa29: more to less ancestral) with a coloured bar indicating geographic region of individual (see Fig. 1). Asterisk (*) indicates a potential recent second-generation European backcross. Mean confidences (assignment posterior probabilities) are provided below chromosomes. Tajima’s D was calculated across each chromosome using 500KB windows of SNPs on **(E)** Ssa01, **(F)** Ssa23, **(G)** Ssa08, and **(H)** Ssa29 for different ancestry groups in North America and groups combined. Within outlier regions (highlighted in gray), arrows indicate significant selection (upward arrow = balancing selection; downward arrow = positive selection) with arrow colour corresponding to group experiencing significant selection.

For Ssa08/Ssa29, individuals were split into three ‘pure’ ancestry groups: European (Q > 0.9 from Europe), North American ancestral (Q > 0.9 from North America) and North American derived (Q < 0.1 from North America) and ancestry was assigned to North American individuals with intermediate Q-values (0.1 < Q < 0.9; n = 256). Within outlier regions, ancestry was primarily assigned as either North American type (ancestral or derived) with limited evidence of European introgression (Fig. 6CD). Tajima’s D was consistent with balancing selection acting on the ancestral North American type within the outlier regions of Ssa08 (W = 613, p = 0.004; Fig. 6G) and Ssa29 (W = 748, p = 5.91e-05; Fig. 6H). Evidence of balancing selection was also found in these regions when ancestral and derived North American types were analyzed together (p values < 0.0003) but not for the derived type alone (Fig. 6GH). No significant evidence of a positive selective sweep was found within outlier windows on Ssa08 or Ssa29 for derived or ancestral North American individuals (p values > 0.234) (see Supplement Fig. S7).

## Discussion

Pleistocene glaciations and interglacial periods drove range contractions and expansions that have greatly influenced the genomes of many northern species (1, 3, 11–13). As glaciers retreated ~10K YBP, several marine species exhibited post-glacial range expansions from Europe into North America (12) and signatures of historical transAtlantic gene flow have been documented in species, such as Atlantic cod (*Gadus morhua*) (36), North Atlantic wolfish (*Anarhichas* spp.) (37), and sea star (*Asterius rubens*) (38). In Atlantic salmon, colonization of formerly glaciated regions of North America likely involved individuals from both western and eastern Atlantic refugia (16), leading to secondary contact in some regions. Our study identified novel widespread variation in North American Atlantic salmon associated with a chromosomal translocation (Ssa01/Ssa23) reflecting evidence of trans-Atlantic secondary contact as revealed by high levels of heterogeneous European ancestry (>50%) on these chromosomes, contrasting low levels genome-wide (~3%). Our study also found novel and extensive variation associated with a chromosome fusion (Ssa08/Ssa29) likely representing a derived polymorphism for the fusion that evolved within the North American lineage prior to the post-glacial range expansion.

Spatial genetic structure of Ssa01/Ssa23 outliers (but not Ssa08/Ssa29) showed a latitudinal clinal pattern that was consistent with geographic distribution of European-mitochondrial DNA evidence in North America (16, 21, 23, 39), providing evidence of post-glacial secondary contact in northern regions such as Labrador and parts of Newfoundland (NL). The capacity for trans-oceanic migration coupled with changing environment (ice sheet retreat) likely enabled trans-Atlantic secondary contact in these northern regions as salmon from both refugia competed to colonize newly available habitats. Indeed, European salmon still undergo large-scale ocean migrations to major feeding grounds in these northern regions (40). Tagging studies have documented the migration of salmon from Norway, Ireland, and the United Kingdom to the Labrador Sea off West Greenland (41) where the salmon fishery is ~20% European-origin, and has historically been >50% (42, 43). Further, consistent with our results, previous evidence revealed an increasing frequency of European ancestry from west-to-east in southern NL with a transition at the Burin Peninsula (21). Although our NL sites are west of this break, the nearest site (Garnish; GAR) showed greater membership to the European cluster than other NL sites. Our observation of secondary contact in Labrador also agree with reports of European mitochondrial haplotypes in the region (16, 21), yet our study is the first to demonstrate how widespread this contact likely was across systems. These signatures of secondary contact are not consistent with contemporary human mediated movement of salmon (i.e., stocking and aquaculture) (16), and generally, individuals in our study appear to show clear evidence of European ancestry in only portions of their genome.

While trans-Atlantic secondary contact is the most parsimonious explanation for variation in the Ssa01/Ssa23 translocation, spatial genetic patterns for Ssa08/Ssa29 were inconsistent with secondary contact and the ‘European-type’ genetic cluster predominated in many North American rivers although further analyses revealed differences within the ‘European-type’ cluster between continents. Our data suggest that the ancestral North American karyotype for Ssa08/Ssa29 is ‘European-like’ (*i.e.*, no fusion) and, interestingly, more similar to Europe than the derived North American type (*i.e.*, fusion). The derived Ssa08/Ssa29 fusion polymorphism likely arose within the North American lineage prior to post-glacial colonization and has not reached fixation, consistent with our finding of balancing selection maintaining this polymorphism. Although evidence of recent secondary contact could also be present within the Ssa08/Ssa29 region, we found limited European ancestry within the region. It is likely that during recent secondary contact recombination could freely occur between the ancestral North American and introduced European karyotypes (*i.e.*, both no fusion for Ssa08/Ssa29) allowing the breakdown of LD in the region.

Although Ssa01/Ssa23 and Ssa08/29 outlier blocks may originate from different histories, it is likely that these regions include important adaptive variation within populations and this is consistent with the observed signatures positive selection on Ssa01/23 and balancing selection on Ssa08/29. Indeed, quantitative trait loci (QTLs) for life history variation, including age at physiological adaptation to seawater (smoltification) and sexual maturation, have been found on several chromosomes including Ssa01, Ssa08, and Ssa23 using trans-Atlantic backcrosses, where Ssa23 in particular showed genome-wide significance for early smolting (44). Ecologically relevant QTLs across six salmonid species were found to be significantly enriched on Ssa01 (and syntenic regions in other species), and several species showed QTLs for morphological, physiological, and life history traits (45) within the outlier block identified in our study. Additionally, many SNPs on Ssa01, Ssa08, and Ssa29 explained regional genetic structure associated with differing thermal regimes in Labrador (46), thus we predict that these genomic regions harbour many genes of ecological significance in Atlantic salmon.

An important implication of our study is the identification of extensive karyotype variation within and among individuals and populations of North American Atlantic salmon. Interestingly, previous limited karyotyping of North American salmon (primarily derived from St. John River, NB) revealed that individuals were heterozygous for the Ssa08/Ssa29 fusion but fixed for the Ssa01/Ssa23 translocation (19), consistent with our findings in the same geographic region. Individual heterozygosity for the Ssa26/28 fusion was also found in the previous study (19), but our study did not detect outlier blocks associated with this rearrangement potentially indicating the Ssa26/28 fusion has not reached significant LD thresholds or that we have not sampled geographic regions where fusion variation is present. Nonetheless, variation in all three rearrangements suggests that chromosome number (2N) could range between 54 and 58 in North America, consistent with karyotyping studies reporting this range of population-specific mode 2N with evidence of inter- and intra-population variability (19, 47–50). Within individuals, it may be possible for rearranged chromosomes to pair and recombine with crossing over highly restricted at the fusion and fission points thus leading to high LD blocks detected here that are further strengthened by patterns of recombination in male salmonids that are highly localized towards telomeric regions (51). Further, residual tetraploidy coupled with generally low recombination rates of salmonids (52) may protect against the possible negative consequences of karyotypic differences given that intraspecific variation in chromosome number is especially common in salmonid species (53). Homeologous regions identified for Atlantic salmon chromosomes (51) could help compensate for potential costs in hetero-karyotypes. Alternatively, gene flow may be restricted between individuals with differing karyotypes via sexual selection processes (*i.e.*, pre- and/or post-copulatory mate choice), which are often employed by salmonids to promote genetic quality of their offspring (54–57).

In conclusion, we report the first evidence of widespread variation in major chromosomal rearrangements in North American Atlantic salmon that were previously expected to lead to incompatibilities between the eastern and western Atlantic populations (20). Our results suggest that variation in chromosomal rearrangement tracks different evolutionary histories in Atlantic salmon, where the Ssa01/Ssa23 translocation shows evidence of substantial heterogeneous introgression of European DNA into North America likely reflecting evidence of post-glacial trans-Atlantic secondary contact. Conversely, widespread differences associated with a chromosomal fusion (Ssa08/Ssa29) likely represents variation that evolved within the North American lineage, suggesting that the prevalent ancestral North American karyotype is ‘European-like’. Although our study did not directly karyotype individuals and European sampling was limited to Norway, our work indicates that some genomic regions are not suitable for trans-Atlantic discrimination, and this has important management implications for mixed-stock fisheries assessments (40, 42, 58) and quantifying the impacts of wild and farmed (*i.e.*, introduced European broodstock) salmon interactions (59). Our study highlights how admixture events during post-glacial range expansions and genomic architecture can facilitate large-scale intraspecific genomic diversity that can drive evolutionary change which may have contributed to the colonization success of Atlantic salmon in North America.

## Materials and Methods

### Sample collection and genotyping

Atlantic salmon from 80 populations spanning Norway and Canada (Fig. 1 and Table S1) were genotyped using a 220,000 target, bi-allelic SNP Affymetrix Axiom array developed for Atlantic salmon by the Centre for Integrative Genetics (CIGENE, Ås, Norway). Sample collection, preparation, and genotyping for Norwegian sites (n = 54) and Canadian sites in Labrador (n = 17) were previously described in Barson, *et al.* (60) and Sylvester, *et al.* (61), respectively. Samples from other Canadian sites in New Brunswick (n = 2 sites), Nova Scotia (n = 3), and Newfoundland (n = 4) were collected for previous studies (62, 63) and genotyped using the same protocol as Sylvester, *et al.* (61). In total, our analyses included 546 and 913 salmon from Canada (salmon parr) and Norway (salmon adults), respectively.

### Detection of individual-based differences in genomic architecture

The R package *pcadapt* (32) was used to detect genomic regions associated with individual-based differences in genomic architecture across 26 North American salmon populations. Minor allele frequency (MAF) cutoff was set to 0.05 resulting in a total of 104,119 SNPs used in the analysis. We tested multiple values of K (number of principal components; PC) ranging from 1 to 50. We expected that as more PC axes were retained in the analysis, we would increasingly detect outlier SNPs representative of interindividual variation while accounting for population level variation. The final number of PC axes retained (K = 40) was determined by visual inspection of the scree plot. The R package *qvalue* (64) was used to transform p-values for all SNPs into q-values to control for false discovery rate (65). The generated q-values were plotted using the Manhattan plot function in the R package *qqman* (66). From the Manhattan plot, chromosomes characterized by regions with large numbers of loci with significant *q*-values (*i.e.*, large outlier blocks) were identified and further examined with subsequent analyses.

### Linkage disequilibrium of outlier blocks

Linkage disequilibrium (LD) was calculated among all outlier SNPs (*q*-value < 0.05) on chromosomes containing large outlier blocks. Salmon populations from Canada and Norway were analyzed separately. Pairwise LD (R^2^) values were determined using PLINK (67) and visualized using heatmap.2 function in *gplots* (68). In the case that high LD regions were found between outliers on two separate chromosomes, which may occur for regions involved in structural rearrangements, pairwise LD values between all loci (with MAF > 0.05) on the two chromosomes were also visualized in the same way as described above. High LD regions on chromosomes were further examined by plotting mean pairwise LD across the chromosome for each SNP (MAF > 0.05).

### Population and individual level genetic structure of outlier blocks

Spatial genetic structure was examined for outlier blocks. For each outlier block, we selected all significant SNPs (*q*-value < 0.05) within the block region. Significant SNPs from outlier blocks were combined and analyzed together if high LD was detected between regions from different chromosomes that have a known rearrangement. We combined these SNPs because our genomic positions are based on the European Atlantic salmon genome, and thus positions will differ for these chromosomes in North American salmon.

First, Bayesian clustering analysis was performed on datasets of outlier blocks using the program STRUCTURE v2.3.4 (33) where runs were implemented through the R package *parallelstructure* (69). Three independent Markov Chain Monte Carlo (MCMC) runs using 100,000 burn-in and 500,000 iterations were performed for each value of K (*i.e.*, genetic clusters) ranging from 1 to 15. We determined the optimal number of K by calculating the ΔK statistic (34) using STRUCTURE HARVESTER (70). Using the best K, admixture coefficients from STRUCTURE were used to calculate the proportion of population-specific membership to each cluster.

Population-specific allele frequency patterns across latitude were examined for the outlier datasets for Canadian populations with the Norwegian allele considered as the major allele for each locus. Allele frequencies were modelled across latitude using generalized linear models.

Next, we examined individual-based differences, where we compared individual genetic relationships for different outlier blocks. In addition to outlier blocks, we chose a panel of 1500 randomly selected non-outlier SNPs (*q*-value > 0.05 and excluding chromosomes with outlier blocks) to reflect genome-wide neutral population structure. For the outlier and non-outlier datasets, we compared genetic distances among individual from Europe and North America using a neighbor-joining (NJ) tree based on Cavalli-Sforza and Edwards (71) chord distances calculated in POPULATIONS v1.2.33 (72) with 1,000 bootstrap replicates on loci. FigTree v1.4 (73) was used to visualize the genetic relationships. Genetic structure was also examined using principal component analysis (PCA) in the R package *adegenet* (74). All individuals were used for the PCA. For the NJ tree, all North American individuals were included but only a subset of representative Norwegian individuals were chosen from five arbitrarily selected populations across different geographic regions (Enningdalselva, Nausta, Årgårdsvassdraget, Elvegårdselva, and Risfjordvassdraget). This subset of individual was sufficient to reveal a clear trans Atlantic split for genome-wide markers and was consistent with the PCA using all individuals (see Results).

### Ancestry and selection along chromosomes

PCAdmix (35) was used to identify ancestry along each chromosome. For each chromosome, genotypes were phased using BEAGLE 3.0 (75). Given that histories and thus relationships may differ for different outlier blocks, PCAdmix was used to determine ancestry across all chromosomes based on the results Bayesian clustering for outlier datasets separately. For the identified Ssa01/Ssa23 outlier dataset, individuals were split based on their co-ancestry coefficients (STRUCTURE Q-values) where pure Europe and pure North American groups included European individuals with Q-value < 0.1 (n = 628) and North American individuals with Q-value > 0.9 (n = 355), respectively. All individuals with some evidence of potential European introgression (Q-value < 0.9; n = 191) were assigned ancestry along each homologous chromosome. For Ssa08/Ssa29, we assumed three potential ancestry types (see Results): European (Q value < 0.1 from Norway; n = 381), North American ancestral (Q value < 0.1 from Canada; n = 101) and North American derived (Q value > 0.9 from Canada; n = 189). All individuals with intermediate Q values (0.1 < Q value < 0.9; n = 250) were assigned ancestry. Ancestry along each chromosome was assigned to 20 SNP windows and ancestry assignments were associated with a posterior probability. Posterior probabilities of each assignment were averaged for each SNP window to assess relative confidence in assignment. SNPs were excluded from the analysis if they were monomorphic across all pure samples.

In addition to ancestry, PopGenome (76) was used to examine selection on outlier regions. Individuals within North America from the extremes of the Bayesian clustering assignments (Q values < 0.1 or Q values > 0.9) were used for the analysis. For Ssa01/Ssa23, individuals were grouped as North American (Q > 0.9) or North American-European admixed (Q < 0.1) which corresponded to only individuals with strong evidence of European introgression. For Ssa08/Ssa29, individuals were grouped as North American ancestral (Q < 0.1) and North American derived (Q > 0.9) and this was guided by results of NJ tree. Groups were analyzed separately and together for calculations of Tajima’s D across chromosomes in 500KB windows. Statistical significance of positive and balancing selection within outlier regions were tested against chromosome-wide values using a Mann-Whitney U test. Last, we used the Sweepfinder composite likelihood method in SweeD (77, 78) on each chromosome to detect 500KB windows with signatures of positive selection. We ran SweeD analysis separately for the same groups used for Tajima’s D calculations. We categorized regions with composite likelihood ratios greater than the 95% quantile of chromosome-wide values as under selection in all comparisons. Statistical significance of selective sweeps within outlier regions were also tested against chromosome-wide values using a Mann-Whitney U test.

## Acknowledgements

We thank the Centre for Integrative Genetics (CIGENE) for genotyping the range-wide data used in the present study.

## Data Accessibility

Data will be made available pending manuscript acceptance.

## Author Contributions

SJL, IRB, TK, JBH, and PB contributed to the conception and design of the study. SJL, JBH, and TK performed statistical analyses. MC, PB, JBH, and SL provided molecular data and metadata for the study. SJL drafted the manuscript and all authors contributed to the writing and approved the final draft of the manuscript.

